# Dynamically regulated transcription factors are encoded by highly unstable mRNAs in the *Drosophila* larval brain

**DOI:** 10.1101/2022.12.03.518875

**Authors:** Mary Kay Thompson, Arianna Ceccarelli, David Ish-Horowicz, Ilan Davis

**Affiliations:** Department of Biochemistry, University of Oxford, United Kingdom; Mathematical Institute, University of Oxford, United Kingdom

## Abstract

The level of each RNA species depends on the balance between its rates of production and decay. Although previous studies have measured RNA decay across the genome in tissue culture and single-celled organisms, few experiments have been performed in intact complex tissues and organs. It is therefore unclear whether the determinants of RNA decay found in cultured cells are preserved in an intact tissue, and whether they differ between neighboring cell types and are regulated during development. To address these questions, we measured RNA synthesis and decay rates genome wide via metabolic labeling of whole cultured *Drosophila* larval brains using 4-thiouridine. Our analysis revealed that decay rates span a range of more than 100-fold, and that RNA stability is linked to gene function, with mRNAs encoding transcription factors being much less stable than mRNAs involved in core metabolic functions. Surprisingly, among transcription factor mRNAs there was a clear demarcation between more widely used transcription factors and those that are expressed only transiently during development. mRNAs encoding transient transcription factors are among the least stable in the brain. These mRNAs are characterized by epigenetic silencing in most cell types, as shown by their enrichment with the histone modification H3K27me3. Our data suggests the presence of an mRNA destabilizing mechanism targeted to these transiently expressed transcription factors to allow their levels to be regulated rapidly with high precision. Our study also demonstrates a general method for measuring mRNA transcription and decay rates in intact organs or tissues, offering insights into the role of mRNA stability in the regulation of complex developmental programs.

## Introduction

Development of a multicellular organism requires intricate control of protein production in response to specific signals. Much of this control is exercised at the level of transcription, with signaling cascades leading to binding of transcription factors to recognition sites in the genome. Transcription factor binding precipitates production of a target mRNA, and eventually, translation of protein required in a specific cell at a specific time. Another class of gene regulation, known as post-transcriptional regulation, operates on the mRNA after its production. mRNAs can be regulated post-transcriptionally by several means – translation, intracellular localization, and decay, among others. The stability of an mRNA determines whether it is available for translation into protein and whether it reaches the region of the cell where it is required. Regulation of mRNA stability is a key component of gene regulation because the inappropriate presence of even a single molecule of mRNA could have drastic consequences. According to some estimates, one molecule of mRNA directs the synthesis of 6,000 molecules of protein on average^1^. In combination with translational control, rapid degradation of unneeded RNAs can prevent the accumulation of proteins in the incorrect cell type or at the incorrect developmental stage.

Multicellular development involves many rapid transitions in cell types and finely-tuned spatial boundaries. Such cell-fate changes are often accompanied by rapid changes in gene expression. For example, reiterated, segmental organisation of the early Drosophila embryo depends on the striped expression of pair-rule segmentation genes in alternate segment-wide stripes^2^. Expression of these transcripts is highly dynamic^3,4^, with the mRNAs of *fushi tarazu* (*ftz*) and other pair-rule transcript having half-lives of only 7 min^5,6^. Instability of Ftz protein appears also to be critical for correct patterning^7^.

Later in development, the embryo undergoes even more complicated patterning to create tissues and organs, the most intricate of which is probably the brain. In *Drosophila*, neural identities are determined by a defined series of temporal transcription factors^8^, and similar transcriptional cascades have been described in mammalian neural stem cells^9^. For several switches, a rapid transition in expression of successive factors depends transcriptional repression of an early factor by its successor^10^. However, it seems likely that robust switching between stem cell identities also depends on rapid mRNA degradation or translational repression.

Current measurements of RNA stability have largely been derived from single-celled organisms or cultured mammalian cells, and so do not directly address *in vivo* mechanisms of cell-state changes in developing tissues. Very few studies have estimated RNA stability in multicellular organisms^11,12^, and we are unaware of any data describing RNA stability across an entire brain. In this paper, we estimate synthesis and decay rates in the *Drosophila* larval brain genome wide by analysing the metabolic labeling kinetics of more than 7,000 RNAs. We show that decay rates of different RNA species vary by orders of magnitude, with rates differing greatly across different gene functions. In particular, we find that mRNAs that encode transcription factors are particularly unstable, especially if their expression is restricted to a limited number of cell types. Many of these cell-type-specific transcription factor RNAs are derived from genes which targets of histone H3K27 trimethylation, a chromatin modification that is associated with transcriptional repression. Our results suggest a link between transcriptional and post-transcriptional repression during brain development.

## Results

To study RNA dynamics genome-wide, we first devised a protocol to address the technical challenges of estimating decay rates from small tissue samples (see Methods). We found that RNA transcription and decay rates can be estimated from these samples using metabolic labeling, the steady-state assumption, and a single timepoint design^13^. In this method, synthesis rates are estimated directly from a pulse of 4-thiouridine RNA labeling, and decay rates are inferred by comparing the synthesis rates to the measured total RNA levels from the same sample. The steady-state assumption requires that total RNA levels do not change over the course of the assay. To evaluate this assumption, we collected RNA from brains either directly after dissection or after 60 minutes of incubation in our culture conditions. We found that the global pattern of RNA levels was highly correlated between the two time points (r^2^ = 0.94), supporting our use of the steady-state model (Fig. S1A). A similar result was found when we compared brains incubated for 60 min in the presence of 4-thiouridine to brains harvested directly after dissection (Fig. S1B), suggesting that incubation in 500 µM 4-thiouridine does not cause large gene expression changes over this time frame.

We next generated genome-wide estimates of RNA synthesis and decay rates in the larval brain. We added 4-thiouridine (4sU) to L3 larval brains cultured briefly *ex vivo*^14^, and then isolated RNA from the brains after 20 minutes of labeling time (Fig 1A). This time frame is short enough to resolve transcriptional dynamics for many RNAs^15^, yet long enough to allow precision in the the labeling duration and to recover a reasonable amount of labeled RNA. RNAs containing 4-thiouridine were purified and RNA sequencing libraries were constructed from both the labeled 4sU^+^ RNA and the total RNA. 4sU^+^ RNA was rich in unspliced intronic regions relative to total RNA, as expected, because 4sU^+^ RNAs are either recently transcribed or in the process of transcription (Fig. 1B). The data resulting from these sequencing experiments was then used to estimate RNA synthesis and decay rates^13^ and to explore the roles of RNA dynamics in the brain.

**Fig 1:**
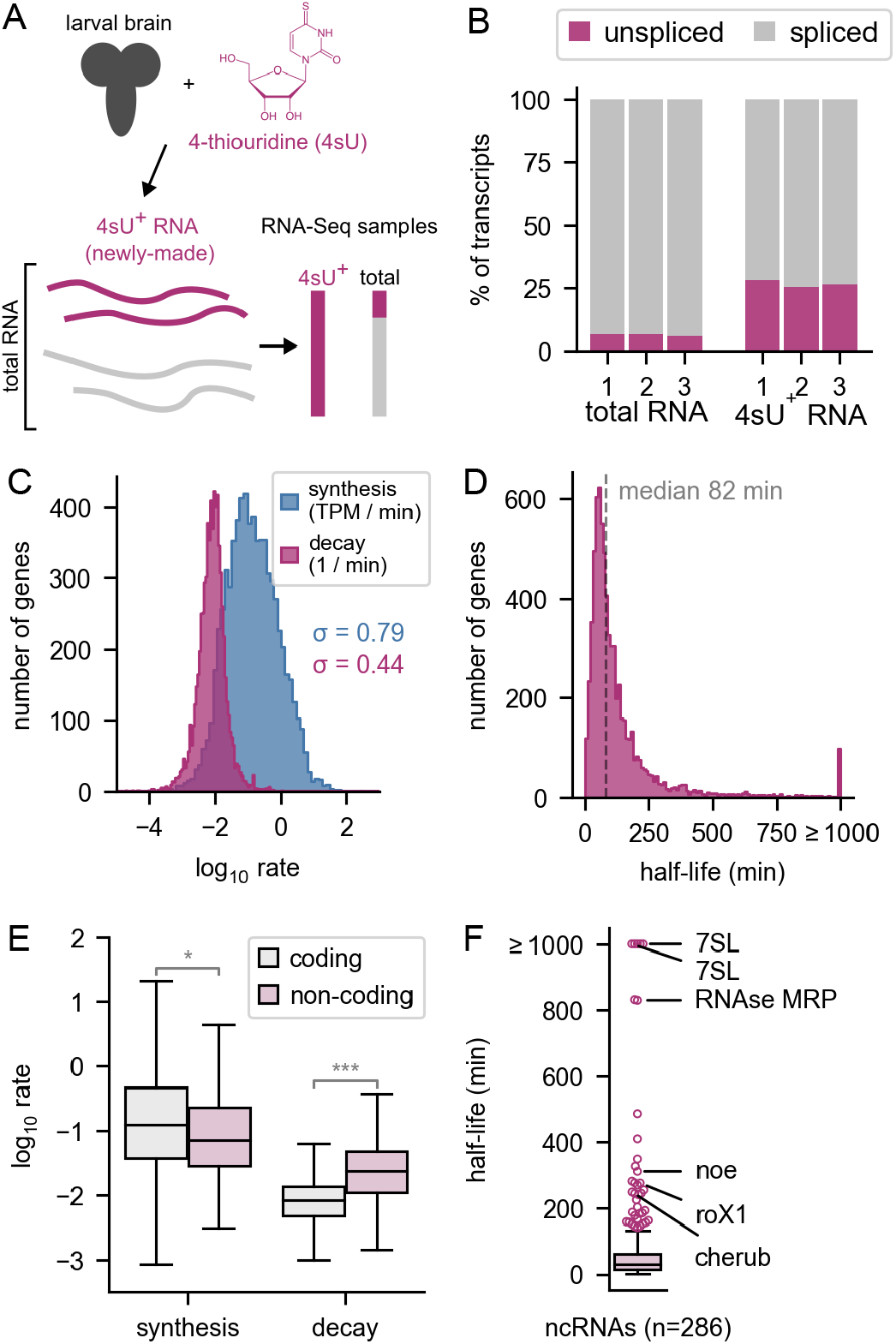
Measurement of RNA transcription and decay rates in larval brains. A) A protocol for measurement of RNA transcription and decay rates in larval brains. B) Estimation of the percentage of spliced and unspliced transcripts in the total and 4-thiouridine enriched (4sU^+^) libraries. C) Histogram of estimated synthesis and C D decay rates among all genes. The standard deviation (s) is used to describe the variation between rates across all genes. The rates were calculated in units TPM / min (transcripts per million per min) and 1 / min. For synthesis rates, s = 0.79 and for decay s = 0.44 for log_10_-transformed rates. D) Half-lives of all RNAs in the larval brain. Half-life estimates were capped at 1000 min, due to estimation uncertainty of very E F long half-lives (see Methods). E) Box plot showing the synthesis and decay rates of coding mRNAs compared to non-coding RNAs (*p<0.05, ***p<10e^-40^). F) Half-lives of non-coding RNAs. Named non-coding RNAs with long half-lives are indicated.

We first used our genome-wide dataset to assess the contribution of transcription and decay rates to the total RNA levels in the brain. Synthesis rates span a wider range than decay rates (σ = 0.79 vs. σ = 0.44 for the log-transformed rates) (Fig. 1C). We transformed the decay rates to half-lives and found that the median RNA half-life was 82 minutes (Fig. 1D), similar to recent estimates in mammalian cells^1,15^. However, the distribution is highly skewed with many short half-lives and a long tail of more stable RNAs. Using our method, we were not able to resolve half-lives greater than approximately 1000 minutes (see Methods). Therefore, we grouped genes with half-lives greater than 1000 minutes into a single bin labeled ≥ 1000 minutes (Fig. 1D). Non-coding RNAs had much higher decay rates than coding transcripts, but only slightly lower synthesis rates (Fig. 1E). Of note, we found that many highly stable non-coding RNAs have known functions – for example, *7SL* and *RNAse MRP* have established metabolic roles, and *roX1* and *cherub* have important developmental functions (Fig. 1F)^16,17^. We then used this dataset as a starting point to identify features linked to RNA stability in the developing brain.

We hypothesized that RNAs which are localized to neurites need to be stable in order to survive their long transit from the nucleus to the periphery. To address this hypothesis, we defined neurite-localized mRNAs based on homology to mammalian mRNAs which were found to be enriched in neurites across several studies ^18^ and compared their decay rates to those of other RNAs. We found that neurite-localized mRNAs have lower decay rates than other mRNAs in the genome (Fig. S2).

They also have higher synthesis rates and higher total RNA levels. We conclude that mRNAs which are present at the distal periphery of neuronal cytoplasmic projects are more highly synthesised, stable, and have higher total RNA levels. We propose that the low decay rates of these mRNAs helps them survive their long journey to the tips of neurites.

To investigate the relationship between the function of mRNAs and their stability, we grouped genes into functional categories using GO Slim annotations and determined the stability profiles of the gene groups (Fig. 2). We found the general principle that mRNA stability was intimately linked to gene function. Genes encoding transcription factors produced RNAs with much lower stability than average (median around 28^th^ percentile). In contrast, genes encoding structural molecules, many of which are ribosomal protein genes, had very high stability (median around 79^th^ percentile). In between these two outliers, gene groups had less severe bias, but many were nonetheless intriguing. RNAs encoding receptors had low median stability whereas RNAs encoding binders of receptors had a bimodal distribution, with some having relatively low and others relatively high stability (Fig. There is only a weak global correlation between RNA decay rate and total RNA level (r^2^ = 0.04), and the correlation is absent when examining the genes encoding transcription factors alone (r^2^ = 0.00) (Fig. S3A). Some highly expressed mRNAs are also unstable. For example, many unstable mRNAs encoding receptors are also highly expressed (Fig. S3B). Together these results argue for tight regulation of RNA stability and coupling of RNA degradation to the functional roles of the mRNA.

**Fig 2:**
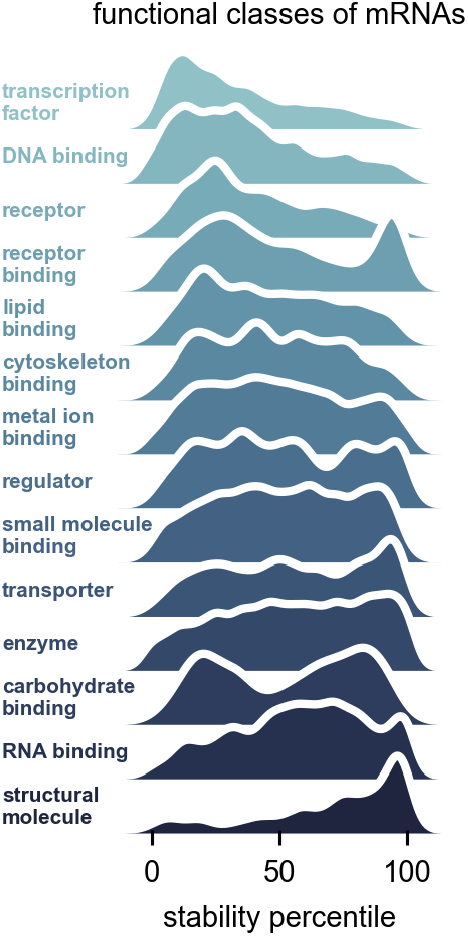
Different functional classes of mRNAs have distinct decay rates. Stability percentile of GO slim molecular function categories for mRNAs. The GO slim categories were taken from the Flybase functional ribbon annotations. The histograms are displayed as kernel density estimates.

Brain tissue consists of many different cell types. We tested whether RNAs that are highly regulated and expressed in only certain cell types would be more unstable than other RNAs that are more widely expressed in the brain. We found that in general, this is not true, but it is true for the subclass of RNAs that encode transcription factors. To classify RNAs as cell type specific or non-specific, we made use of a single-cell RNA-seq study of the larval brain from a similar developmental timeframe (48 hours after larval hatching, early L3)^19^. We classified RNAs as cell type specific (CTS) if they were both statistically enriched in a given cell type and at least 2-fold more abundant in that cell type relative to the rest of the brain. RNAs enriched in certain cell types such as neuroepithelial cells were generally unstable (Fig. 3A). In contrast, RNAs enriched in hemocytes, glia, and several classes of mature neurons tended to be stable. When viewed as a group, CTS RNAs are more stable than average, if RNAs encoding transcription factors are excluded (Fig. 3B).

**Fig 3:**
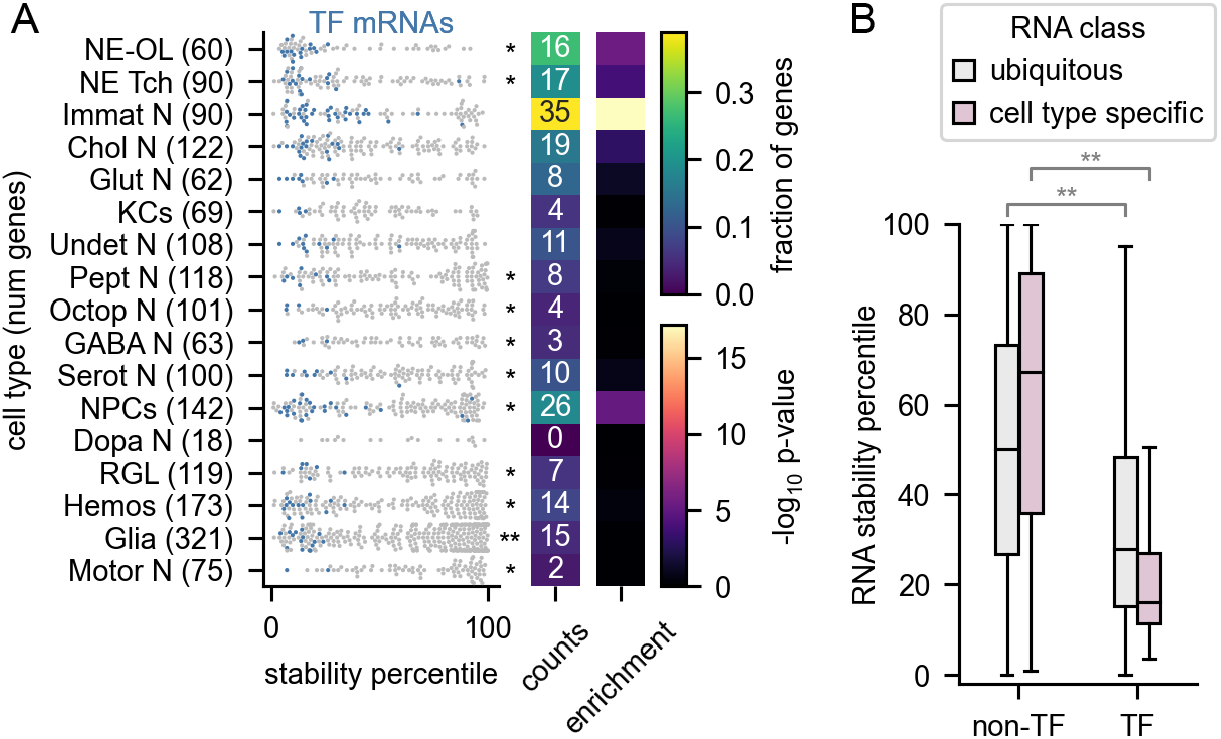
Cell-type-specific transcription factor RNAs are rapidly degraded. A) The stabilities of RNAs (including both mRNAs and ncRNAs) that are enriched in specific cell types are shown. mRNAs encoding transcription factors (TFs) are indicated in blue. The asterisks to the right of the plot indicate that the stabilities for the indicated group of cell-type-specific RNAs are different than other RNAs in the dataset (Mann-Whitney U test). The heatmaps on the right display the fraction of each category that is composed of TFs and the significance of the enrichment. The enriched RNAs were extracted from a single-cell RNA-seq dataset derived from larval brains^19^. Cell type key: NE-OL (neuroepithelia/optic lobe), NE Tch (neuroepithelia/trachea), Glut N (glutamatergic neurons), Immat N (immature neurons), Chol N (cholinergic neurons), Undet N (undetermined neurons), KCs (Keyon cells), Pept N (peptidergic neurons), Octop N (octopaminergic neurons), Serot N (serotoninergic neurons), GABA N (Gabaergic neurons), NPCs (neural progenitor cells), Dopa N (dopaminergic neurons), RGL (ring gland), Hemos (hemocytes), Motor N (motor neurons). B) Stability of cell-type-specific RNAs vs. non-specific RNAs for transcription factor genes (TFs) and other genes (non-TFs). (*p<0.05, **p<10e^-10^, ***p<10e^-40^).

To explore this effect further, we analyzed the functions of RNAs in the top and bottom 10% of stability for CTS RNAs. Stable CTS RNAs were enriched for metabolic processes and regulation of transporters (Fig. S4). In contrast, unstable CTS RNAs were most enriched for transcriptional regulators. Strikingly, CTS RNAs that encode transcription factors were more unstable than other transcription factor RNAs (16^th^ percentile of stability vs. 28^th^ percentile of stability) (Fig. 3B). Among the various cell types, we found that RNAs specific to immature neurons, neural progenitor cells, neuroepithelia, and cholinergic neurons were enriched with mRNAs encoding transcription factors (Fig. 3A).

To put our results into context, we examined the RNA stability and cell type enrichment for another class of regulators, the RNA-binding proteins (RBPs) and the subclass of these which are known to bind mRNAs, the mRBPs. RBPs and mRBPs were only enriched in RNAs specific to neural progenitor cells (Fig. S5A). Cell-type-specific mRBP mRNAs, but not cell-type-specific RBP mRNAs, have lower stability than other cell-type-specific mRNAs (Fig. S5B). These analyses show that cell type specificity is linked to RNA stability, but only for certain classes of genes.

We next asked what features could explain the low stability of cell-type-specific transcription factor RNAs. These RNAs had no significant enrichment for known RNA-binding protein motifs (see Methods)^20^. However, we found that they are targets of epigenetic silencing. Silenced genes are marked by regions of histone H3K27 trimethylation, which often extend through both the gene body and its regulatory regions. These genes are often referred to as Polycomb group (PcG) target genes ^21,22^. To identify potential PcG target genes, we examined H3K27 trimethylation sites in the fly genome collected from L3 larvae as part of the modENCODE project^23^. 339 genes for which we have determined RNA decay rates correspond to genes that have methylation sites within 1 kb of their transcription start site (upstream), within 1 kb of their transcription stop site (downstream) and within the gene body, the region corresponding to the transcript itself (including both introns and exons of the gene). Because these genes are predicted PcG targets, rather than experimentally confirmed targets, we refer to them as extended H3K27me3 genes. These genes have exceptionally low stability when compared to genes without H3K27 trimethylation, and other classes of genes with methylation covering only some of these regions also had lower stability than average (Fig. 4). This observation suggests a link between transcriptional silencing and high RNA decay rates.

**Fig 4:**
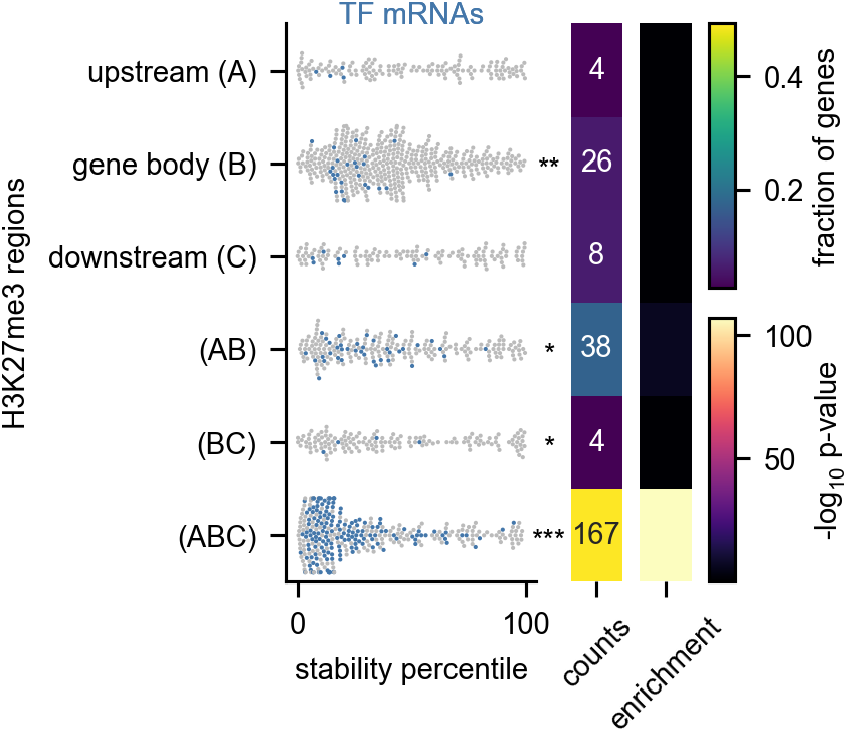
Unstable transcription factor RNAs are targets of H3K27 trimethylation. The RNA stability percentile of genes with H3K27me3 binding sites is shown. The asterisks to the right of the plot indicate that the stabilities for the indicated group of cell-type-specific RNAs are different than other RNAs in the dataset (Mann-Whitney U test). The heatmaps on the right display the fraction of each category which is composed of TFs and the significance of the enrichment. The H3K27me3 binding sites were derived from ChIP-seq of whole L3 larvae which were collected as part of the ModENCODE project^52^. Genes were classed as having a binding site within 1 kb of their transcription start site (upstream (A)), within 1 kb of their transcription termination site (downstream (C)), or within the gene body itself, including both introns and exons (gene body (B)). We assigned genes to a methylation category based on the combination of regions which had a methylation binding site. Each gene is assigned to only one category. We excluded the upstream-downstream (AC) category because of scarcity; only 10 genes fell into this category. (*p<0.05, **p<10e^-10^, ***p<10e^-40^).

We wondered whether the relationships between RNA decay rates and cell type specificity, H3K27 trimethylation, and transcription factor status could be disentangled from each other. These gene groups are highly overlapping, with RNAs encoding transcription factors making up 49% of extended H3K27me3 genes and cell-type-specific transcription factors making up 13% of extended H3K27me3 genes (Fig. 5A). These overlaps are highly significant. Transcription factor genes are enriched for extended H3K27me3 (odds ratio 18.3, CI = 14.4 – 23.2, p < 0.001) and CTS transcription factor RNAs are also enriched for extended H3K27me3 (odds ratio 17.8, CI = 11.8 – 26.9, p < 0.001) (Fig. 5A). The overlap among these gene groups complicates any simple interpretations that do not consider the relationship between them.

**Fig 5:**
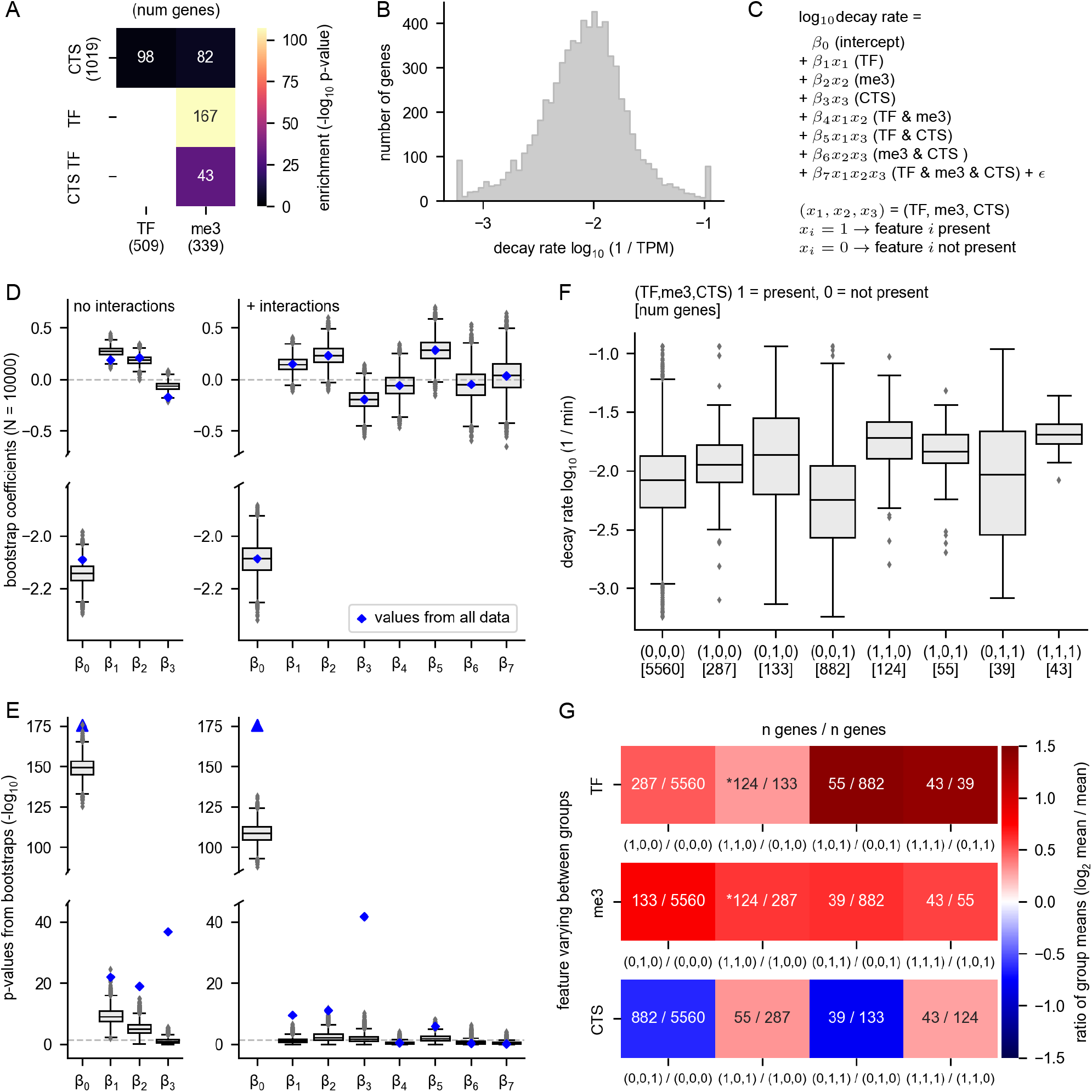
Relationships between RNA decay and gene attributes. The relationship between RNA decay rates and gene membership in the following categories is shown: transcription factor (TF), extended H3K27me3 genes (me3), and cell-type-specific genes (CTS). H3K27me3 target genes were defined as those which have H3K27me3 binding sites upstream, downstream, and covering the gene body, as specified in Figure 4. A) The overlap between each gene group is shown. CTS TFs are cell-type-specific transcription factor genes. The p-value’for the overlap enrichment is shown. Note that the overlap between CTS TF & TF and the overlap between TF & TF is not shown because these categories have 100% overlap by definition. The numbers in parentheses indicate the number of genes in each group. B) The log transformed and winsorized decay rates are shown. C) The equation for the linear regression model is shown. The model predicts the logw decay rate using gene membership in each of the three groups (TF, me3, and CTS) as features. The model was built either with or without interactions between the features. b_0_ − b_3_ indicate the coefficients, with only b_0_ − b_7_ determined for the model without interactions. The model without interactions can be derived from this equation simply by setting the interaction coefficients (b_4_ − b_7_) to 0. D) A multiple linear regression model was built to predict RNA decay rate as a function of TF, CTS, and H3K27me3 status. The blue diamonds indicate the values determined using all data, whereas the grey box plots show the range of values obtained from subsampling the dataset with bootstrapping. Bootstrapping was performed by subsampling each non-overlapping gene group with a unique combination of factors using a sample size equivalent to the smallest group size (n=39). 10,000 bootstrap samples were taken. (Note: the eight non-overlapping gene groups are shown in (F)). E) P-values obtained from the linear regression model in (D). The p-values are the -log_10_ value of the original p-values. The p-values corresponding to b_0_ using the entire dataset are 0, thus the -log_10_ value is undefined, but it could be interpreted as an infinitely large number, and is indicated by the blue triangles. F) The decay rates of all eight possible non-overlapping groups derived from three possible explanatory features is shown. Each group is indicated by a vector (TF, me3, CTS), where 1 = present and 0 = not present. G) For each of the three possible explanatory features (TF, me3, CTS), the mean change between a group which is positive for the feature is compared to the group that is negative for the feature but is identical with respect to the other features. The overlaid numbers indicate the number of genes in each group. An asterisk marks the comparisons which show the effect of adding either TF or me3 to a group which already has the other feature (i.e. adding TF to a group which is already positive for me3 or vice versa).

To determine which features(s) have the largest correlation with RNA decay rates, we performed multiple linear regression to model the decay rates as a function of gene membership in each of the three groups (TF = encodes a transcription factor, me3 = extended H3K27 trimethylation, and CTS = cell type specificity). We found that these features differ in their effects, and that interactions between the features change the interpretation of the effects. Before regression, we log transformed the decay rates and winsorized the top and bottom 1% of the data to reduce the effect of outliers (Fig. 5B). Next, we modeled the decay rates as a function of RNA membership in each gene group, either with or without interactions between the features (Fig. 5C – E). The obtained coefficients indicate the strength and directionality of the effect, whereas the p-values indicate the probability that the coefficient is zero. The coefficients β_0_, β_1_, β_2_, β_3_ are present in both models, but they can differ. For example, β_1_ from the non-interaction model indicates the effect of TF without considering that the TF category overlaps the CTS category, but the model with interactions takes this overlap into account, with the coefficient β_5_ describing the effect of TF and CTS combined. The model with interactions shows that TF has an effect both independently of other categories and in combination with CTS (both β_1_ and β_5_ are positive, Fig. 5D, blue diamonds).

In order to focus our analysis on genes which belong to our groups of interest, we rebuilt the model with 10,000 bootstrapped data sets sampled from our data (Fig. 5D – E). This procedure will give each of our gene sets of interests the same importance in the model, rather than importance proportional to their frequency in the original dataset. For the bootstrapped models, we randomly sampled each independent gene group (groups shown in Fig. 5F) at the same depth (39 genes per group per iteration, which corresponds to the minimum size of all gene groups). The bootstrapped models largely agree with the models built using the whole dataset, and the range of values obtained in the bootstrapped models reinforce confidence in the effects of some features. The coefficients obtained from bootstrapping differ slightly from the coefficients derived from the whole dataset in the model without interactions (Fig. 5D), showing, for example, that the effect of β_1_ (TF) is higher when the analysis is focused on our gene groups of interest than when using the whole dataset. In contrast, the coefficients are nearly identical between the bootstrapped models and the model considering the whole dataset in the case of the model with interactions – this is likely because many of the independent gene groups that are used to determine the interaction coefficients are small and highly or completely sampled in the bootstrapping procedure. We found that β_2_ (me3) and β_5_ (TF & CTS) are positive in nearly all the bootstrapped models, reinforcing our confidence that both H3K27 trimethylation and encoding a transcription factor in combination with cell type specificity are associated with higher RNA decay rates (Fig. 5D, right). β_1_ is positive and β_3_ is negative in most bootstrapped models, showing that encoding a transcription factor and cell type specificity are associated with high and low RNA decay rates, respectively (Fig. 5D, right). Overall, our linear models suggest that both H3K27 trimethylation and encoding a TF are predictors of high RNA decay rates, as is the combination of the features cell-type-specific and encoding a transcription factor.

To address the same question in a different way, we examined the decay rates of each independent gene group in which the combination of features is constant (Fig. 5F). We found that extended H3K27 trimethylation is associated with a similar increase in decay rates regardless of the other features it is combined with, whereas the other groups show more variable behavior. Each group is denoted by a vector which specifies its combination of features, with zero indicating that the feature is not present and one indicating that the feature is present (see Legend). For example, (1,0,0) identifies genes which encode transcription factors, but which do not have any other features. This gene set does not overlap with the set denoted by (1,1,0), which specifies genes which both encode TFs and have extended H3K27 trimethylation. Direct comparison of each independent group gives similar insights to those obtained via linear regression. The CTS feature alone is not a consistent indicator of high RNA decay rate and in some comparisons is associated with a lower decay rate (Fig. 5G). In contrast, extended H3K27 trimethylation is a consistent indicator of high decay rate across all comparisons, whereas the effect of encoding a transcription factor is strongest in combination with the CTS feature (Fig. 5G).

Because of the strong and consistent correlation of H3K27 trimethylation with high RNA decay rates, we wondered whether there might be a functional link between PcG targeting and RNA decay. Such a link would ensure that transiently expressed RNAs could not wreak havoc in another cell type as a consequence of leaky transcription and that they could be cleared effectively before transitioning to another cell state. The most obvious explanation for the link is that convergent evolutionary selection processes have led to both PcG-driven transcriptional silencing and low RNA stability for these genes. However, the most parsimonious explanation for the link would be that the two are functionally related – i.e. whatever processes serve to transcriptionally silence the gene also cause the locus-derived RNA to be rapidly degraded. To distinguish between these possibilities, we examined a dataset which measured both nascent and total RNA levels after knockdown of components of the Polycomb repressive complex 1 (PRC1) in a *Drosophila* neural-derived cell line BG3^24^. As expected, knockdown of the Ph subunit of PRC1 increased the levels of many RNAs at the nascent RNA level. These RNAs also increased in total RNA levels (Fig. S6A – C). Nascent RNA sequencing captures the active transcriptional response to a change. It is expected that a change in transcriptional regulation of a gene would change its nascent RNA levels, but that the total RNA levels would mirror the changes in nascent RNA with a slight time delay depending on the half-life of the RNA in question. If a perturbation acts by altering RNA stability, then we would expect it to change the ratio between the nascent and the total RNA levels. We found that the ratio between the total and nascent RNA levels was slightly increased after Ph knockdown for genes which were defined as probable targets of PcG-driven silencing (PcG domain genes, Fig. S6D). This experiment suggests that Ph depletion primarily derepresses the transcription of PcG domain genes, but that the stability of the resulting RNAs remains similar after they are transcriptionally derepressed. Therefore, we fail to find evidence of a direct causative link between PcG targeting and RNA decay.

Finally, we examined the RNA stabilities for two groups of genes which are known to be spatiotemporally regulated in either the *Drosophila* embryo or larva. One group of genes is involved in neural fate determination, via specifying the developmental timing window in which a neuron is born. Another group of genes regulates anterior-posterior patterning in the *Drosophila* embryo. These groups of genes had low RNA stability, in agreement with our finding that rapid RNA decay is needed for RNAs which are required transiently in development (Fig. 6). We note that five of these genes (*hb, Kr, tll*, and *slp1/2*) are shared between the groups and have functions in both processes. The majority of these RNAs encode transcription factors (Fig. 6). Therefore, we suggest a model wherein mRNAs encoding transiently expressed and dynamically regulated transcription factors are selected to be unstable so that rapid developmental transitions can occur robustly.

**Fig 6:**
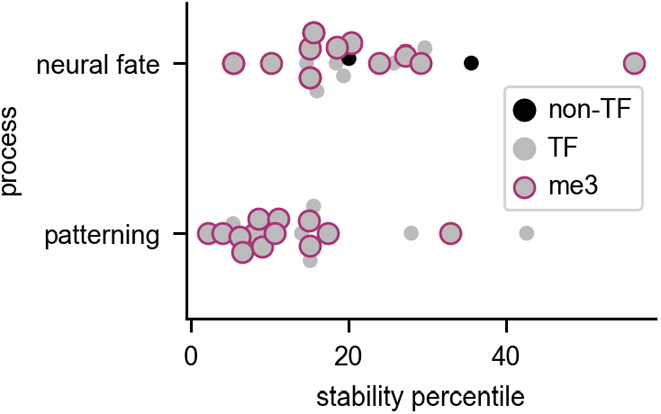
RNAs which are highly regulated during development are rapidly degraded. The stability percentile of RNAs (both mRNAs and ncRNAs) which are annotated as being involved in neural fate determination or early patterning of the *Drosophila* embryo. For neural fate determination, the RNAs are a combination of temporal transcription factor genes and other genes that undergo temporal regulation in neural precursor cells^53^. For pattern formation, the genes were taken from a curated list hosted on the Interactive Fly website and encompassing gene sets known as the gap genes and the pair-rule genes^54^. Most of these genes are TFs; the non-TFs are indicated. Genes which are in our group of extended H3K27me3 genes (me3) are also indicated.

## Discussion

Here, we report a systematic analysis of RNA transcription and decay rates in the developing *Drosophila* brain. We accomplished this feat by adapting 4sU metabolic labeling and sequencing to small tissue samples. Although we collected RNA from 150 larval brains per replicate, we have shown that the experiment can be performed with as little as 5 µg of total RNA which can be collected from as few as 25 brains (see Methods). Our study serves as proof of principle that RNA labelling experiments and kinetic data can be collected from complex tissue types, which are often more biologically relevant than monolayer cells in culture. There are many potential uses for our dataset. For example, our analysis of decay rates identified many lncRNAs with higher than average RNA stability. Among these lncRNAs was the RNA *cherub* (half-life = 248 min), which was recently found to have an important function in brain tumor formation^17^. It will be interesting to examine the functions of other high stability lncRNAs, which are likely to have important roles.

Using this data, we showed that transcription rates span a larger range than decay rates, but decay rates are nonetheless biologically relevant. As shown in previous studies of RNA decay from tissue culture cells and single-celled organisms, RNA decay is tightly coupled to the function of the encoded RNA^1,15,25,26^. Interestingly, not all high expression RNAs are stable. For example, some RNAs encoding receptors are highly abundant but also unstable. Such dynamics might allow cells to respond to changes in conditions rapidly, even for abundant RNAs.

Perhaps the most striking finding was the high turnover rate of RNAs encoding transcription factors. Although some past studies of RNA kinetics in tissue culture cells have observed lower stability for RNAs encoding transcription factors, our study takes this observation a step further. We found that the low stability status of these RNAs is linked to their tendency to be transiently expressed and developmentally regulated. These characteristics of the unstable transcription factor RNAs are demonstrated via their enrichment in certain transient cell populations, such as among immature neurons. Likewise, their high levels of H3K27 trimethylation in data derived from whole larvae indicates that these genes tend to be kept in a transcriptionally silent state in most cell types.

There are two possible models to explain the relationship between transcriptional regulation and the high decay rates of these RNAs. The first possibility is that the transient TF RNAs contain cis-acting elements, such as RNA-binding protein motifs. On the whole, we have not found evidence for universal RBP targeting of TF mRNAs, but it is possible that evolutionary pressures have selected sequence characteristics that destabilize these RNAs on a case-by-case basis and/or that destabilization depends on complex combinatorial action of multiple RNA-binding proteins that cannot be easily detected.

A second possibility is that transcriptional regulation of these RNAs is directly linked to their high degradation rates. Using data from another study^24^, we found that knockdown of PRC1 subunit Ph generally increases both the nascent and total RNA levels of PcG target RNAs, which argues against a direct effect of the PRC1 complex on RNA decay. However, this observation does not rule out the possibility that PcG targeting could be linked to RNA degradation of the same gene product through an unknown mechanism. This RNA destabilizing mechanism might remain active even after the gene is derepressed via experimental manipulation or *in vivo* during development. There is a growing body of evidence that factors which regulate the RNA life cycle can be deposited on the RNA during the process of transcription, a process which has been shown to depend on the promoter^27–31^. Likewise it seems possible that factors which regulate transcriptional silencing could also recruit RNA destabilizing factors which could be deposited on an RNA during transcription. An exciting candidate for a PRC linked destabilizing factor is the rixosome, a complex involved in rRNA processing that was recently shown to interact with PRC complexes^32^. Future experiments involving careful manipulation of the PRC complexes and ablation of interactions with candidate RNA decay regulators will be needed to determine if a causative link between PcG targeting and RNA decay exists.

## Materials and Methods

### Choice of labeling approach and protocol optimization

We considered different approaches for estimating RNA decay rates from our small tissue samples. We decided to employ a pulse labeling approach with 4-thiouridine (4sU) and to extract RNA kinetic parameters from a single timepoint^13,33^. Classically, RNA decay has been measured with a pulse-chase approach. However, this method is difficult to perform with tissue samples. A pulse period and subsequent chase period in culture will expose the tissue to more time and potential damage from the *ex vivo* conditions. Furthermore, many chase timepoints are needed to capture the half-lives of RNAs at different ends of the half-life spectrum. In addition to the extra resources need to process these numerous samples and their replicates, taking timepoints from tissue involves destructive sampling – that is they are collected from different sets of animals. Calculations which involve comparison between timepoints will compound error for measurement of RNA half-lives from noisier sources such as tissue. In the single timepoint method, both labeled RNA and total RNA are quantified from the same timepoint, and the resulting data is used to estimate RNA synthesis and decay rates using a steady-state model^13,33^.

### *Drosophila* culture and RNA extraction

*Drosophila melanogaster* wild type strain Oregon-R flies were raised on standard cornmeal-agar medium at 25°C. For each replicate 150 wandering third instar larvae were collected and their brains were dissected in brain culture media (BCM: 80% v/v Schneider’s *Drosophila* medium (Thermo Fisher 21720024), 20% v/v heat inactivated fetal bovine serum (Sigma F4135) FBS, and 10 µg/ml human insulin (Sigma I9278). The BCM used here is based on a previous recipe used for longterm culturing of larval brains^14^ but modified to exclude whole larval extract, which contains RNAses and non-brain RNAs that could contaminate our sequencing libraries. Brains were collected in small batches of 15-20 in order to minimize total dissection and preparation time before labeling, which was kept to under 20 minutes. Brains were transferred to 1.7 ml epitubes and the media was replaced with BCM containing 500 µM 4-thiouridine (4sU, Carbosynth NT06186) for 20 minutes. For the incubation condition testing experiments shown in Figure S1, 50 brains were collected per replicate and the brains were incubated for 60 minutes. Our current version of the labeling protocol is able to recover adequate 4sU-labeled RNA from as few as 25 brains, corresponding to ∼5 µg of total RNA as input for the 4sU RNA purification. In addition to the methods described below, we now recommend using SPRI bead-based purification instead of chloroform extraction and using buffers which further increase 4sU-labeled RNA purification specificity (protocol available on request).

Labeling was performed with tubes laid on their side with gentle shaking in the dark (80 rpm on an orbital shaker, New Brunswick Innova 2000). At the end of the labeling period, the brains were quickly washed with Schneiders media + 500 µM 4sU. All incubation and wash steps used 200 µl of the specified media. The wash with Schneider’s medium was performed to remove traces of FBS, which we found to have high levels of a thiol-reactive molecule that that could possibly interfere with our 4sU RNA purification approach. The samples were then flash frozen in liquid nitrogen. Next, RNA was extracted with the hot acid phenol method. Brains were homogenized on ice in SEA buffer (50 mM NaOAc, pH 5.2, 10 mM EDTA, 0.3% Sarkosyl) for 20 seconds using a motorized pestle. After homogenization, 1% SDS was added, and the extraction was continued as previously described^34^. Previous experiments using Trizol for extraction led to sporadic failure of 4sU purification, which we believe was due to reducing agent carryover from Trizol to the sample.

### Purification of thiolated RNAs and sequencing library construction

RNA was biotinylated in a 360 µl reaction containing 10 mM HEPES 7.5, 1 mM EDTA, and 24 µg of brain RNA. After adding other components, MTSEA biotin-XX (MTS-biotin, Biotium #90066) was added to a final concentration of 0.04 mg/ml as suggested^35^. The stock concentration of MTS-biotin was dissolved in anhydrous DMSO (ThermoFisher, D12345) to a concentration of 0.2 mg/ml and stored at -80 °C before use. The biotinylation reactions were purified with 2 rounds of extraction with an equal volume of chloroform and cleaned up with isopropanol precipitation. Pellets were resuspended in 50 µl of H2O and subjected to purification of on µMACs streptavidin columns as previously described, although reaction volumes were scaled down two-fold (i.e. 50 µl RNA added to 50 µl of beads)^36^. Pulldown RNA was isopropanol precipitated and resuspended in 10 ul H2O. All precipitations were done with 150 mM NaOAc, pH 5.2, 50% isopropanol, and and 50 µg/ml GlycoBlue (ThermFisher, AM9516), then washed in cold 75% ethanol.

Ribosomal RNA depletion was then performed using biotinylated RNAs which are anti-sense to the 5S, 5.8S, 18S, and 28S ribosomal RNAs, as previously described^37^ with adaptations. We used *in vitro* transcribed anti-sense rRNA against the 5S, 5.8S, 18S, and 28S rRNAs (see below). For total RNA samples, 500 ng of RNA was mixed with asrRNA in a 50 µl hybridization reaction. All recovered pulldown RNA was mixed with asrRNA in a 25 µl hybridization reaction. For each 100 ng of input RNA, a mix containing 0.3 pmol of each asrRNA was added per each 100 ng of input RNA, which is approximately 2-fold molar excess over the rRNA in the sample. The input hybridization reaction was purified with 250 µl of Dynabeads MyOne Streptavidin C1 (Thermo Fisher, 65001) in 100 ul binding volume. The pulldown hybridization reaction was purified with 25 µl of beads in 25 µl volume. Beads were washed in Dynabeads solution A prior to use. The binding reactions were incubated 10 min at 22 °C in the thermomixer at 600 rpm. The supernatant (rRNA depleted) was then cleaned up and treated with DNAse using the RNAqueous-Micro Total RNA Isolation Kit (Thermo Fisher, AM1931).

The RNA was eluted in 10 µl from the kit and 6 µl was input into the library construction with the Lexogen SENSE Total RNA-Seq Library Prep Kit for Illumina (Lexogen 009). Libraries were PCR amplified using primers with unique 6mer i7 indices with 19 cycles (input libraries) or 18 cycles (pulldown libraries). Libraries were pooled and sequenced on an Illumina HiSeq 2500 and sequenced as 2 × 125 bp paired-end reads with 2 × 9 cycles of additional index read sequencing. Samples for the incubation condition testing experiments were processed similarly, except that ribosomal RNA subtraction was not performed and 500 ng of total RNA was used to construct poly(A)-primed libraries using the QuantSeq 3′ mRNA-Seq Library Prep Kit FWD for Illumina (Lexogen 015) and amplified for 13 – 15 PCR cycles. Quant-seq libraries were loaded onto an Illumina NextSeq 500/550 High Output v2 cartridge and subjected to single-end sequencing on a NextSeq 500 instrument for 85 cycles with 6 cycles of index sequencing.

### *In vitro* transcription and spike-ins

Drosophila rRNAs corresponding to the 5S, 5.8S, 18S, and 28S were cloned in the anti-sense orientation behind the T7 promoter in the ERCC plasmid backbone and linearized with BamHI. In vitro transcription was performed with the T7 Megascript kit (Thermo Fisher, AM1334) with 40% of the UTP substituted with Biotin-16-UTP (Tebubio N-5005). Reactions were treated with TURBO DNAse (Thermo Fisher, AM2238) and then cleaned up with the RNA Clean & Concentrator-25 kit (Zymo Research, R1017) and eluted with water.

We added two types of synthetic RNA spike-ins to use for quality control. Non-thiolated spike-in RNAs were from the Lexogen SIRV E2 mix (Lexogen 025.03) and thiolated spike-in RNAs (mix m2) were derived from a subset of the ERCC plasmids donated by NIST including: ERCC-00057, ERCC-00073, ERCC-00077, ERCC-00104, ERCC-00112, ERCC-00142, ERCC-00150, ERCC-00162, ERCC-00165, and ERCC-00168. The thiolated spike-ins were made using T7 RNA polymerase (NEB M0251) on BamHI-digested templates. The transcription reactions contained 0.5 mM of each NTPs and additional 4sUTP (TriLink N-1025) added in proportion to the number of Us in each transcript: >350 Us, 0.125 mM; 190-350 Us, 0.25 mM; 100-190 Us, 0.5 mM; <100 Us, 1 mM. The thiolated spike-ins were quantified with the Qubit RNA BR assay (Thermo Fisher, Q10210) and mixed at equal final concentrations. The non-thiolated spike-ins were added to both the input and pulldown samples before ribosomal RNA removal. The thiolated spike-ins were added to the pulldown samples before biotinylation and pulldown, whereas they were added to the input samples before ribosomal RNA removal. For the 20 min labeling experiments, 2.25 ng of thiolated spike-ins were added to each input sample and 0.25 ng were added to each pulldown sample. 4.8 ng of Lexogen E2 mix was added to each input sample and 6.1 ng was added to each pulldown sample. For the 60 min incubation test samples, 0.15 ng of thiolated spike-ins and 0.31 ng of Lexogen E2 mix was added to each total RNA sample.

### RNA-seq data processing

Adapter removal and/or quality trimming were performed following recommendations from Lexogen. For SENSE RNA-seq libraries, the first 9 nt of the R1 read, the first 6 nt of the R2 read, and adapter sequences were removed with Cutadapt^38^ using the command “-m 20 -u 9 -U 6”. For the QuantSeq libraries, BBDuk^39^ was used with the parameters “k=13 ktrim=r mink=5 qtrim=r trimq=10 minlength=20”. Raw sequence data from BG3 cells that was generated using the Ion Total RNA-seq Kit v2 (Thermo Fisher, 4475936)^24^ was downloaded from the SRA and used without further processing because the data had already been trimmed. *Drosophila melanogaster* genomic and transcript sequences were downloaded from Ensembl (release 99, assembly BDGP6.28). Kb-python^40^ was used to build a Kallisto^41^ index with the coding, non-coding, and spike-in transcripts, as well intronic sequences with 30 nt of flanking sequence on either side. Kallisto was used to estimate counts mapping to each intron and each transcript using the options “--rf-stranded” for the SENSE RNA-seq libraries and the options “-l 220 -s 130 --single --fr-stranded --single-overhang” and “-l 100 -s 100 --single --fr-stranded --single-overhang” for the QuantSeq and Ion Total RNA-seq libraries, respectively. Transcript per million (TPM) values were then recalculated for both intronic and exonic regions for each gene after removing reads which were assigned to ribosomal RNA or spike-in RNAs. INSPEcT^13^ was then used to estimate RNA synthesis, processing and decay rates from the 4sU-labeled and total RNA data. Snakemake^42^ and Conda^43^ were used to run the RNA-seq pipeline and manage software dependencies. Downstream analysis and figure construction was completed using open-source scientific computing software, including Numpy, Scipy, Pandas, and Matplotlib, run in Jupyter notebooks^44–48^. Further details of the RNA-seq pipeline and the generated figures are available on Github at https://github.com/marykthompson/foursu_timecourse and https://github.com/marykthompson/brain_stability.

### RNA-Seq data analysis

Before downstream analyses, the data sets were filtered to exclude genes which were not adequately expressed. For each experiment, we chose a cutoff of 10 counts in at least 2/3 of the libraries for at least one condition. Because small RNAs are not recovered quantitatively by general RNA-seq protocols, in our analysis we focus on mRNAs and non-coding RNAs >200 nt (this includes both lncRNAs and other non-coding RNAs such as the signal recognition particle 7SL RNA). For analysis of decay and synthesis rate trends in specific gene groups versus other genes, the Mann-Whitney U-test was used. To calculate the error in the synthesis and decay measurements reported in Table S1, we used the variance of the synthesis and total RNA levels reported by INSPEcT^13^. We converted these variance measures to coefficient of variation (CV) values and also calculated CV values for degradation rates based on the formula used by INSPEcT for decay rate calculation at steady-state (decay rate = synthesis rate/(total RNA – pre-mRNA)). We observed that some genes had very long calculated half-lifes (> 1000 min), but that the variation between half-lives from these genes had large inter-replicate variation due to estimate uncertainty. We chose 1000 min as an upper limit based on the expected purification specificity of 4sU^+^ RNA relative to total RNA in the 4sU^+^ sample. Based on an empirically determined estimate from spike-in RNAs, we determined that some unlabeled RNAs will be present in the 4sU^+^ RNA sample. We chose a cutoff of a 1:100 ratio of 4sU^+^ to mature RNA in the total RNA library. This ratio corresponds to a calculated half-life of 1386 min, which we rounded to 1000. For scatterplots where experiments are compared, a pseudocount corresponding to the minimum value across experiments (i.e. min TPM) was added to all values before log transformation to allow plotting.

### Comparison of RNA stability data with other datasets, gene lists, and gene features

Genes encoding transcription factors were obtained from the Flybase Gene Groups page. RNA-binding proteins were taken as genes mapping to GO term (RNA binding, GO:0003723) and mRNA-binding proteins were taken as genes mapping to GO term (mRNA binding, GO:0003729). Genes enriched in specific cell types in the L3 brain were taken from a single-cell RNA-seq dataset and were defined as genes statistically enriched in a given cell type (p < 0.05) and with a fold-change enrichment of at least 2-fold^19^. PcG domain genes were taken from a study that identified them based on their high enrichment levels in both Ph Chip-seq and H3K27me3 ChIP-chip^24^. Neurite-localized RNAs were extracted from a meta-analysis and correspond to genes found to have significant neurite enrichment (p < 0.1) in at least three studies^18^. DIOPT 9.0^49^ was used to extract *Drosophila* homologs of gene sets from other species. The DIOPT query was run using https://www.flyrnai.org/cgi-bin/DRSC_orthologs.pl and filtered to remove homologs found in fewer than 8 sources (DIOPT score > 7). The search for enriched RBP motifs was done with Transite using the TSMA mode against their entire database of RBP motifs^20^. GO analysis was performed with clusterProfiler^50^ and the returned categories were simplified with the semantic similarity score cutoff set to 0.5. For modeling decay rates as a function of gene features, decay rates were first log_10_ transformed and winsorized at the top and bottom 1%. The rates were fit to a linear model that considered the effect of each feature alone and in combination with each other (see Fig. S6C). The model was run on either the entire dataset or subsampled 10,000 times with bootstrapping. Bootstrapping was performed by subsampling each non-overlapping gene group with a unique combination of features using a sample size equivalent to the smallest group size (n=33). Regression was performed with the OLS function in the Statsmodels package^51^.

### Data Deposition

RNA sequencing data, including processed files summarizing RNA kinetic rates and expression levels, have been deposited in the Gene Expression Omnibus under accession GSE219202.

## Acknowledgements

We thank the US National Institute of Standards and Technology (NIST) for generous donation of the DNA plasmids used for making thiolated spike-in RNAs. We also thank Lexogen for providing the SIRV spike-ins used in this study and for performing part of the sequencing as part of the Lexogen Research Award. Sequencing of some samples was performed by the NGS Facility at the Vienna Biocenter Core Facilities (VBCF), member of Vienna Biocenter (VBC), Austria. Sequencing of other samples was performed in the Oxford Department of Zoology with technical assistance from Amanda Williams.

## Author contributions

M.K.T. and I.D. designed experiments and wrote the manuscript. M.K.T. performed experiments and data analysis. A.C. contributed to data analysis via mathematical modeling.

---

**Fig S1:**
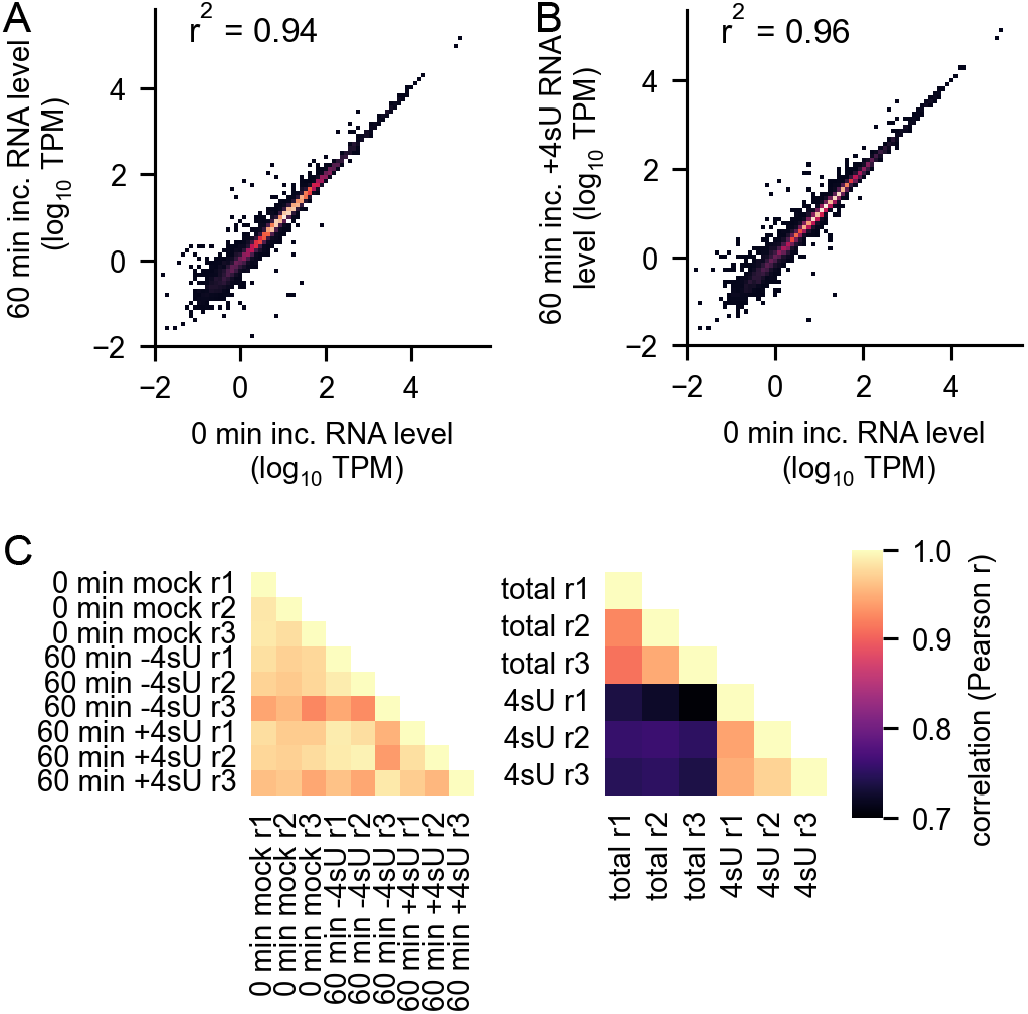
Short-term brain incubation *ex vivo* is compatible with pseudo steady-state assumptions. A) Comparison of total RNA levels from brains harvested directly after dissection to those harvested after incubation in assay media for 60 min. B) Comparison of total RNA levels from brains harvested directly after dissection to those harvested after incubation in assay C media with 500 µM 4sU for 60 min. C) Reproducibility between the RNA-seq samples used in this study. The TPM values were compared for each replicate for all genes passing a cutoff of at least 10 counts in all libraries. For A – C, the Pearson r^2^ value is shown.

**Fig S2:**
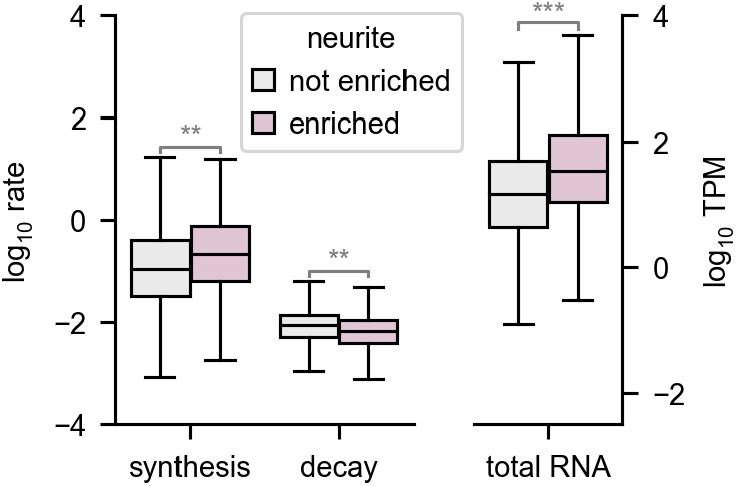
Neuronally localized RNAs have high stability. Boxplot of RNA stability for RNAs which have been identified as enriched in neural projections. Projection localized RNAs were defined as those with enrichment p<0.1 in at least three different studies^18^ (*p<0.05, **p<10e^-10^, ***p<10e^-40^).

**Fig S3:**
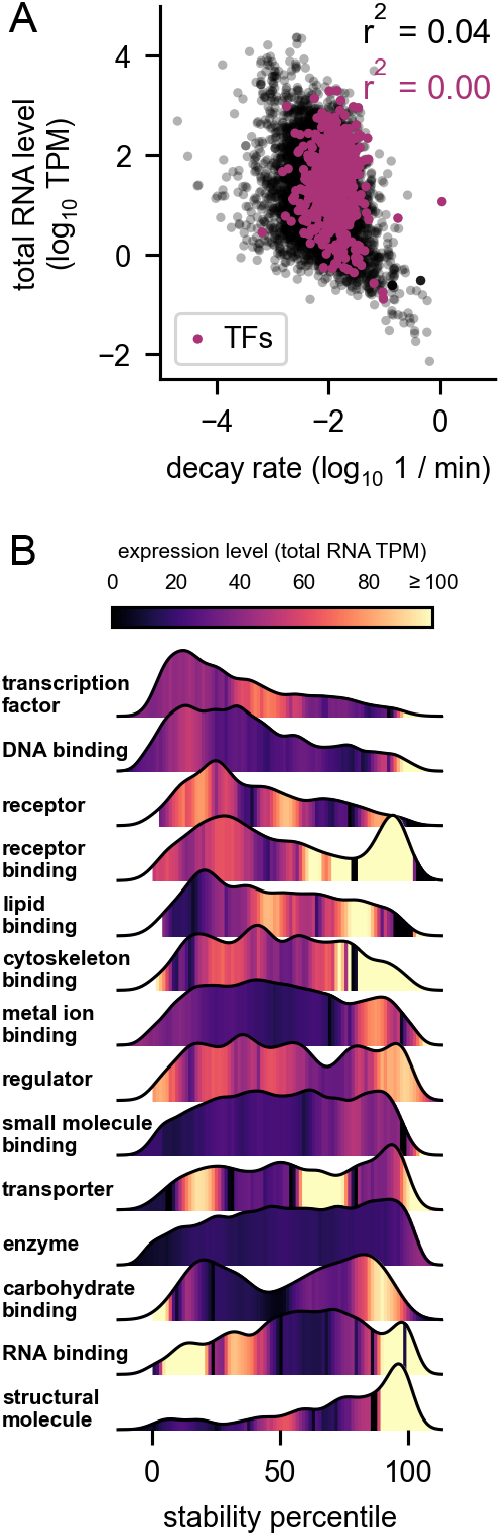
The relationship between decay rates, functional classes, and RNA abundance. A) Relationship between the total RNA level and the decay rate. Spearman’s r^2^ is shown. The axis display limits were chosen to show the main data trend. B) Stability percentile of GO slim molecular function categories. The GO slim categories were taken from the Flybase functional ribbon annotations. The histograms are displayed as kernel density estimates. The median expression level of the RNAs binned by stability percentile in each category is shown with the color map.

**Fig S4:**
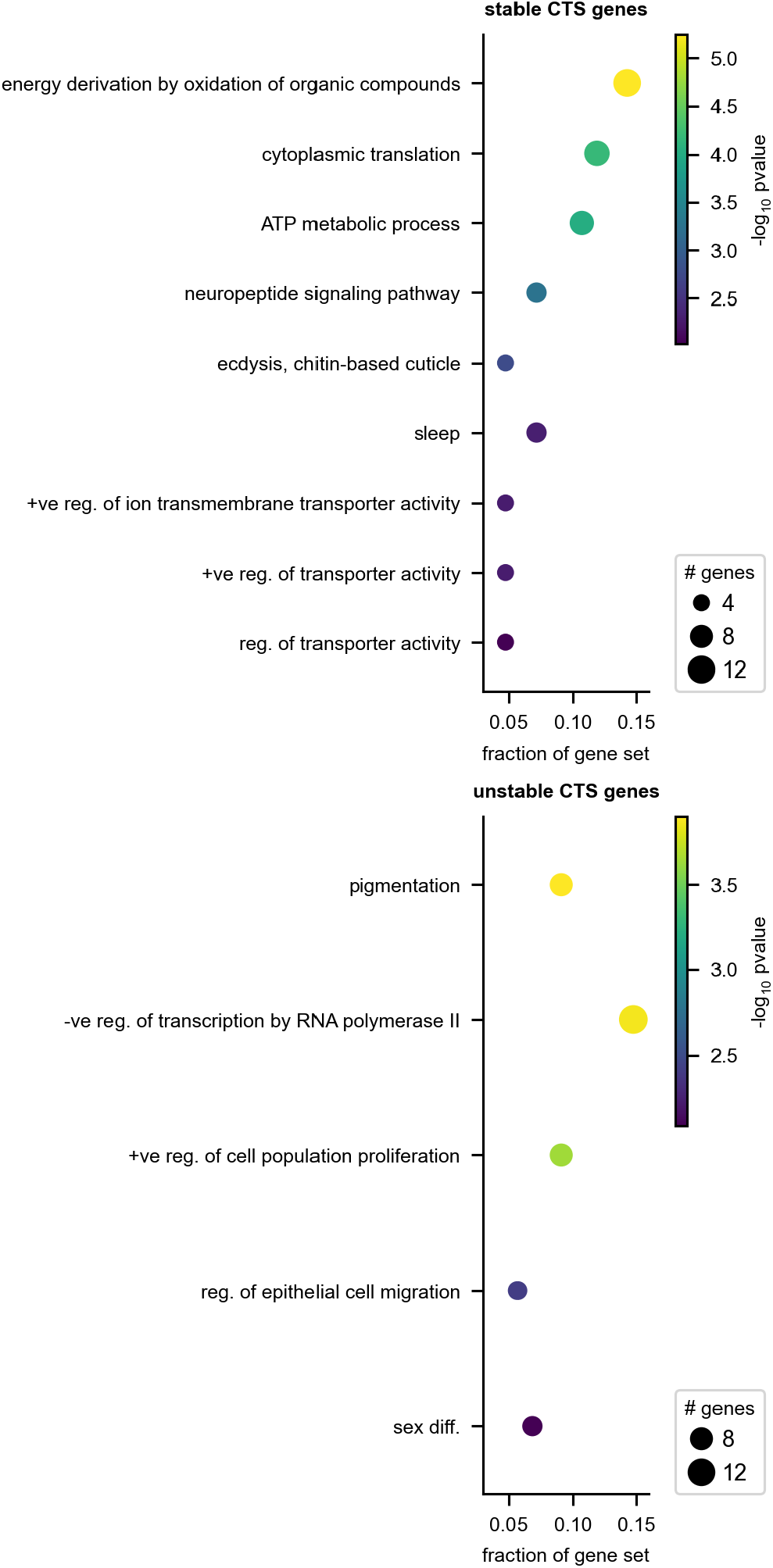
Biological functions of stable and unstable cell-type-specific RNAs. Enriched gene ontology (GO) terms in the biological process namespace is shown for the top 10% most stable cell-type-specific and bottom 10% most unstable cell-type-specific (CTS) RNAs.

**Fig. S5:**
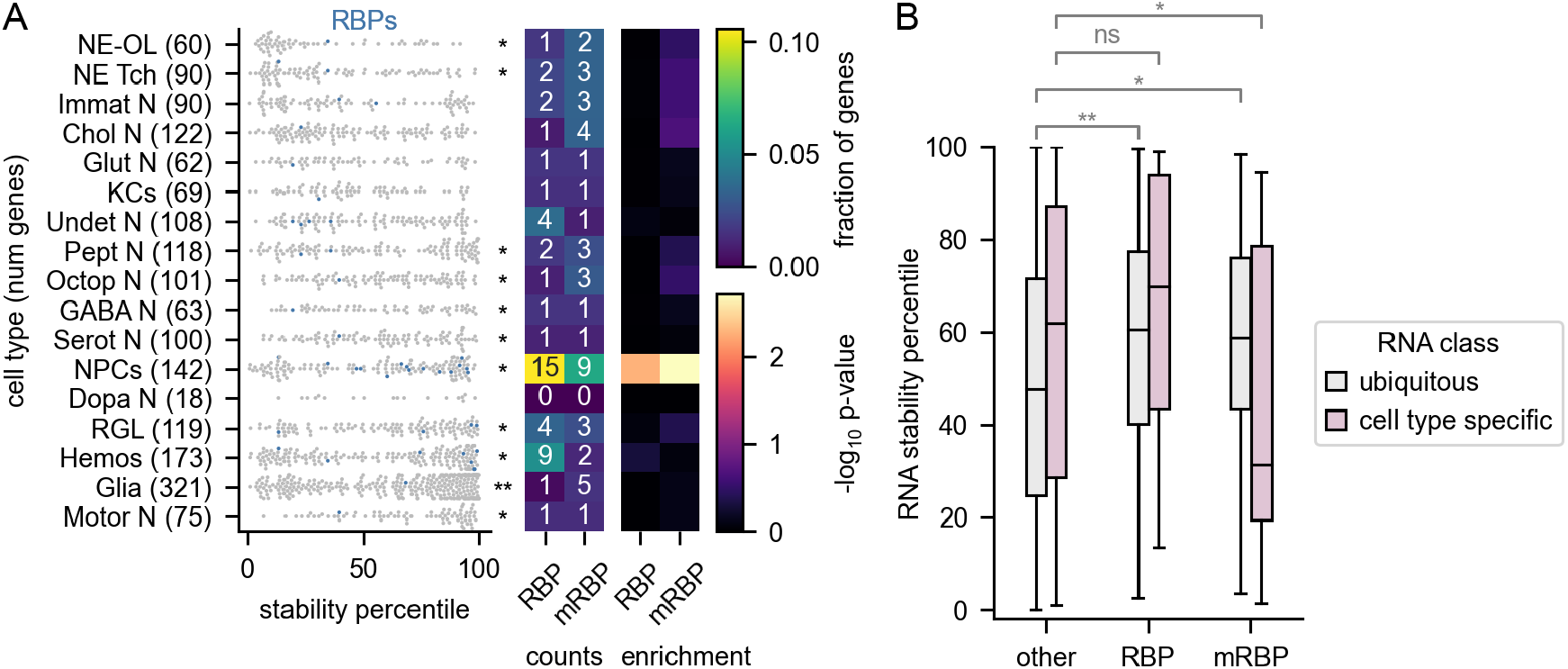
mRNAs encoding RNA binding proteins are enriched in neural progenitor cells. A) The same plot shown in Fig. 3A, but with the mRNAs known to encode RNA-binding proteins (RBPs) highlighted. mRBPs are RBPs which are known to bind mRNA and are a subset of RBPs. B) Stability of cell-type-specific RNAs vs. ubiquitous RNAs for mRNAs encoding RBPs or mRBPs (*p<0.05, **p<10e^-10^, ***p<10e^-40^).

**Fig S6:**
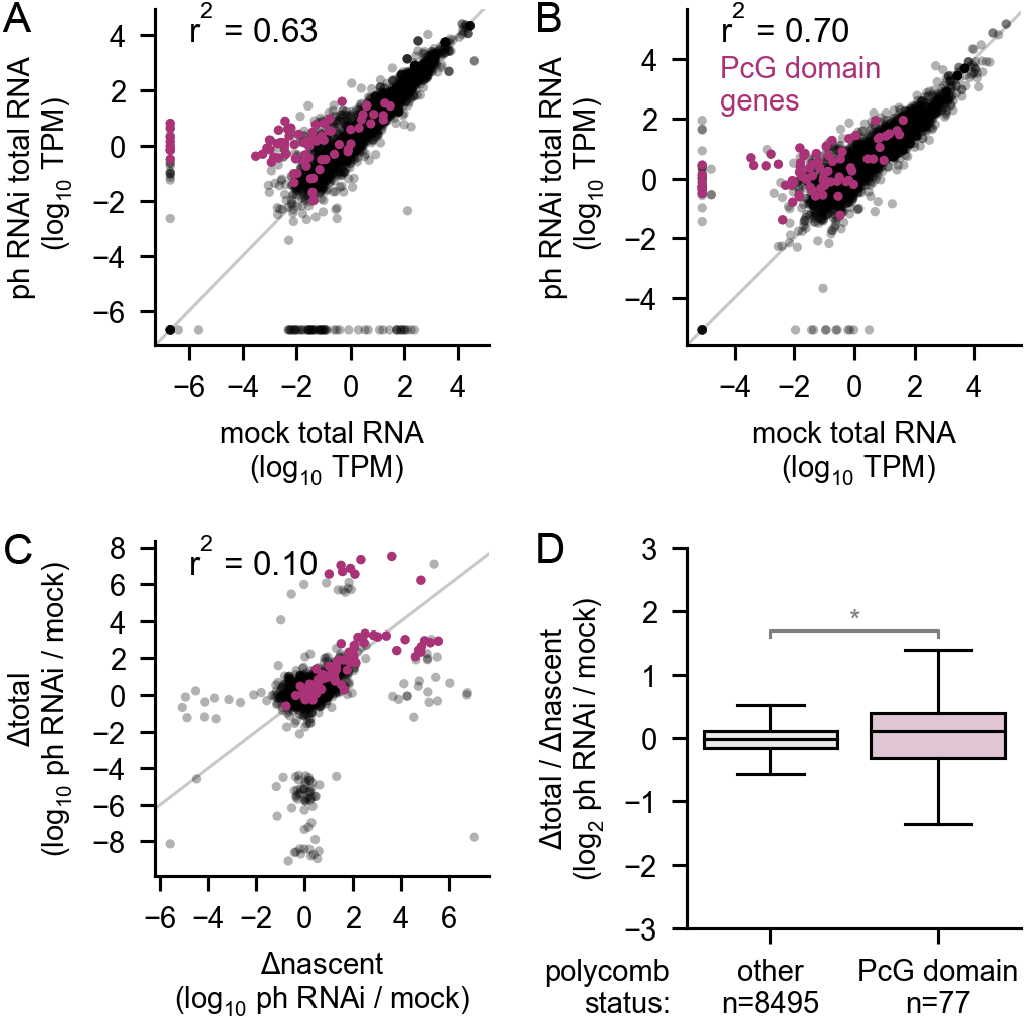
Unstable TF RNAs decrease in nascent and total RNA levels after Polycomb depletion. A – B) The change in total RNA levels (A) or nascent RNA levels (B) after knockdown of Polycomb repressive complex 1 (PRC1 complex) subunit Ph in *Drosophila* neuronal cells (BG3 cells)^24^. The purple dots show genes that the authors defined as PcG domain genes, determined by ChIP-seq against Ph and H3K27me3 in the same study. C) The change in total RNA relative to the change in nascent RNA after Ph knockdown in BG3 cells. D) The ratio of the change in total RNA to the change in nascent RNA for PcG domain genes vs. other genes in BG3 cells.

